# Molecular Surface Descriptors to Predict Antibody Developability

**DOI:** 10.1101/2023.07.18.549448

**Authors:** Eliott Park, Saeed Izadi

## Abstract

Understanding the molecular surface properties of monoclonal antibodies (mAbs) is crucial for determining their function, affinity, and developability. Yet, robust methods to accurately represent the key structural and biophysical features of mAbs on their molecular surface are still limited. Here, we introduce MolDesk, a set of molecular surface descriptors specifically designed for predicting antibody developability characteristics. We assess the performance of these descriptors by directly benchmarking their correlations with an extensive array of *in vitro* and *in vivo* data, including viscosity at high concentration, aggregation, hydrophobic interaction chromatography (HIC), human pharmacokinetic (PK) clearance, Heparin retention time, and polyspecificity. Additionally, we investigate the sensitivity of these surface descriptors to methodological nuances, such as the choice of interior dielectric constant for electrostatic potential calculations, residue-level hydrophobicity scales, initial antibody structure models, and the impact of conformational sampling. Based on our benchmarking analysis, we propose six *in silico* developability rules that leverage these molecular surface descriptors and demonstrate their superior ability to predict the clinical progression of therapeutic antibodies compared to established models like TAP. ^1^

## Introduction

The development of therapeutic antibodies has transformed the field of medicine, offering new and promising treatments for a diverse array of diseases, including cancer, autoimmune disorders, and infectious diseases.^2–5^ To ensure the effectiveness and success of these therapeutic antibodies, several factors come into play, such as their functional properties, binding affinity, and overall developability.^6,7^ One crucial factor that influences these attributes is the molecular surface properties of antibodies. The molecular surface acts as the interface for recognition and binding events, dictating the specificity^8–10^ and affinity^11–13^ of interactions between antibodies and their targets. In addition, the molecular surface of antibody variable (Fv) domains exhibits a wealth of structural and chemical features that can significantly impact their conformational stability,^14,15^ colloidal stability,^7,15–22^ chemical stability,^15,23,24^ and behavior in surface-protein interactions.^25,26^ All of these properties ultimately influence the cost, time of development, and the success of therapeutic antibodies in clinical trials.

The electrostatic surface potential is an important surface property that plays a key role in understanding various biological processes, such as molecular recognition, protein interactions, and properties.^27^ Accurately describing the electrostatics of biomolecules, including antibodies, is challenging due to their longrange nature, the strong dependence on the protonation states of titratable groups, and the complex influence of the surrounding solvent.^28^ Commonly used electrostatics-based descriptors in the context of antibody developability assessment often overlook important factors such as long-range interactions, interior/exterior dielectric environments and solvation effects. For instance, Sharma et al. proposed sequencebased calculations of net charge and charge symmetry parameter (FvCSP) of the Fv to assess antibody viscosity,^29^ neglecting the effects of 3D structure and solvation in the calculations. Agrawal et al.^30^ introduced spatial charge map (SCM) revealing a correlation between larger negatively charged patches on Complementarity-Determining Regions (CDRs) and higher viscosity of antibody solutions. The SCM approach used a structure-based approach and calculated the charge distribution based on the clustering of atomic partial charges for the exposed residues. Similarly, Raybould et al.^1^ introduced Patches of Positive Charge (PPC) and Patches of Negative Charge (PNC) scores, which considered absolute residue-level charge values based on pre-assigned protonation states and took into account the proximity of charged residues. The MOE software also offers descriptors such as ensemble charge of the Fv (ens_charge_Fv) and uses the distribution of atomic partial charges on the molecular surface to define protein charge patches.^31^ While existing methods have shown varying degrees of success in predicting charge-driven developability characteristics, there is a lack of descriptors that specifically quantify the strength and distribution of surface electrostatic potentials (EP) on different antibody domains, which are crucial for selecting and screening humanized candidates.^9,10,25,26^

Hydrophobicity descriptors are of equal importance in assessing the overall function and developability of antibodies,^1,29,31,32^ yet determining their relevance and accuracy in predicting various developability characteristics continues to be a challenge. LogP-based methods such as AggScore^33^ and MOE’s hydrophobicity patches^31^ have been widely used, and have proven valuable in predicting hydrophobic interaction chromatography (HIC) and aggregation.^34,35^ LogP-based scores are particularly critical in profiling small molecule drugs and are fundamental to Lipinski’s Rule of 5.^36^ In addition, several residue-based hydrophobicity scores tailored specifically for antibodies have been proposed, including the Fv Hydrophobicity Index (HI),^29^ Spatial Aggregation Propensity (SAP),^32^ and Protein Surface Hydrophobicity (PSH).^1^ These methods not only differ in their underlying calculation methods but also in the selection of residue-level hydrophobicity scales. For instance, the HI, SAP and PSH methods utilize the Eisenberg,^37^ Black & Mould,^38^ and Kyte-Doolittle^39^ scales, respectively. Additionally, customized hydrophobicity scales have been developed, specifically aimed at predicting developability characteristics such as hydrophobic interaction chromatography (HIC) or reversed-phase high-performance liquid chromatography (RP-HPLC).^40^ The use of diverse hydrophobicity scales in these methods highlights the lack of consensus regarding the significance and optimal choice of residue-level scales for practical predictions of antibody developability. For example, while Raybould et al.^1^ suggested their hydrophobicity score (PSH) is relatively unaffected by the choice of residue-level scale, Franz Waibl et al.^41^ demonstrated that HIC retention times prediction is heavily influenced by the hydrophobicity scale employed. Overall, the choice of hydrophobicity calculation method and the underlying scale can significantly impact the assessment of antibody developability characteristics.^41^

Beyond the impact of underlying methods for surface properties calculations, the accuracy of such descriptors relies heavily on the quality of the predicted antibody structures models.^43^ Recent advancements in deep learning methods specifically designed for antibody structure prediction, such as AB2,^44^ DeepAb,^45^ Equifold,^46^ have significantly improved the accuracy of modeling the CDR loops. However, it remains unclear to what extent the varying accuracy of these models in representing the ground truth X-ray crystal structures translates to actual improvment in the accuracy of structural and surface descriptors directly related to antibody developability.

Furthermore, the impact of generating conformational ensembles, as opposed to using single-static structures, on the outcomes of structure-based computational models for developability predictions is still not fully understood. Antibodies, particularly their CDR loops, undergo continuous transitions between different conformational states, which can affect their structural and developability properties.^47–49^ However, in many quantitative structure–property relationship (QSPR) studies, single-static antibody structures are used in order to achieve computational efficiency. There is currently no consensus on the influence of conformational sampling on predictive accuracy of structure and surface descriptors, with some studies suggesting inconsistent improvement in hydrophobicity prediction,^41^ while others demonstrate its significant effect.^32,50^ Further study is needed to gain a better understanding of the interplay between antibody structure models, conformational ensembles, and the accurate prediction of surface properties for assessing antibody developability.

In this work, we explore alternative approaches to calculating electrostatics and hydrophobicity properties of antibodies. We present a set of surface-based representations specifically tailored for antibodies, and conduct a comprehensive analysis to evaluate the performance and relevance of these surface descriptors by directly benchmarking them against key empirical data such as viscosity, aggregation propensity, hydrophobic interaction chromatography (HIC), and specificity. Furthermore, we investigate how sensitive our surface descriptors are to methodological variations, such as the selection ofcinterior dielectric constants for electrostatic potential calculations, the choice of hydrophobicity scales, the sensitivity to the initial structure models, and the impact of conformational dynamics. Lastly, we propose six *in silico* developability rules using our molecular surface descriptors and demonstrate their superior predictive performance for the clinical progression of therapeutic antibodies when compared to established models like TAP^1^ and MOE’s protein property scores.^31^

## Results

### Mapping the key structural and biophysical features to the molecular surface

Figure 1 illustrates the conceptual framework employed to map the key structural properties of antibodies to their molecular surface. A detailed description of this framework can be found in the Methods section. In brief, we employ the AB2 antibody structure models^44^ to build the initial structure of the variable domain. Unless otherwise stated, we also generate an ensemble of Fv conformations through 5ns Gaussian-accelerated Molecular Dynamics (GaMD) simulations.^52^ A discussion on the sensitivity of our analysis to the choice of the initial structure model and the impact of conformational sampling is provided in subsequent sections.

**Figure 1:**
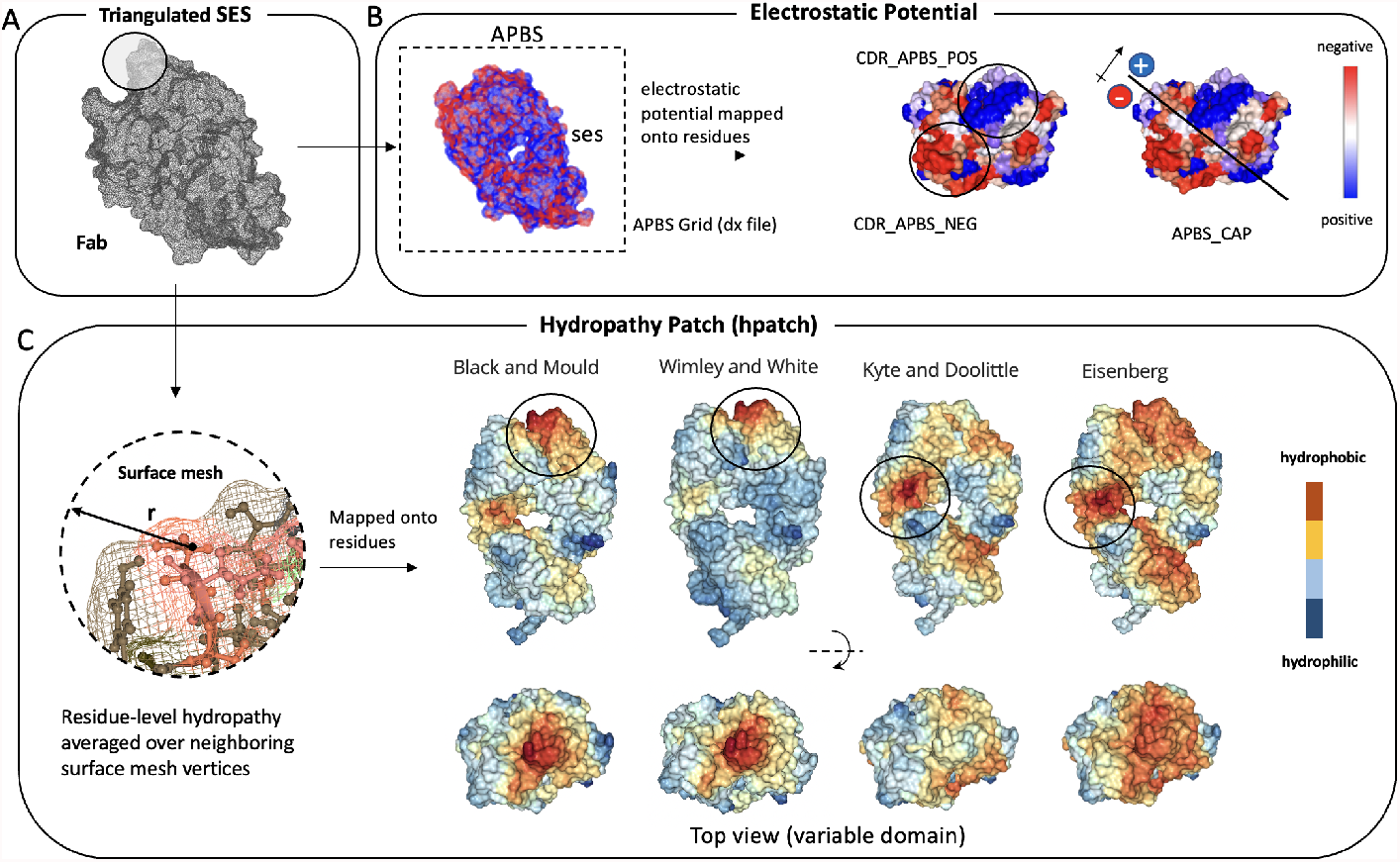
A) Triangulated solvent-excluded molecular surface of a Fab structure generated using Nanoshaper. ^42^ B) Grid-based electrostatic potential from APBS mapped onto the triangulated surface. The potential values are assigned to atoms, and the sum is assigned to corresponding residues. C) Hydrophobicity values assigned to each vertex on the triangulated surface based on residue-level hydrophobicity scales. The values on each vertex are re-assigned by averaging hydrophobicity values from neighboring vertices within a cutoff distance of 10. The largest hydrophobic patch could be on the variable domain (WW and BM) or the elbow (KD and EI), depending on the hydrophobicity scale.

Starting from a single-conformation structure of an antibody Fab or Fv, we generate a triangulated surface mesh (Figure 1A). To determine the electrostatic surface properties, we calculate the continuous electrostatic potential using APBS^53^ and assign the potential values from a discretized cubic grid to the mesh vertices. By summing the positive and negative values across the mesh surface associated with each residue, we calculate the surface electrostatic potential per residue. Subsequently, the integration of surface electrostatic potential over all the residues in the key antibody domains (Fab, Fv, or CDR) provides APBS EP values per each domain (Figure 1B). To capture the asymmetry in the electrostatic potential of the variable domain, we calculate the product of the sum of electrostatic potentials on the VL and VH domains (Figure 1B). This approach mirrors the charge asymmetry propensity highlighted by Sharma et al.^29^ as a crucial factor in antibody self-association.

In contrast to the traditional qualitative approach of visualizing electrostatic potentials using APBS at a single conformation, the method described here provides a quantitative assessment of surface electrostatic potential on a per-residue and per-domain basis. This allows us to calculate average electrostatic potential values per residue or per domain over an ensemble of conformations generated by Molecular Dynamics or other methods. By incorporating dynamical information, our method offers a more rigorous and comprehensive representation of the antibody’s electrostatic properties.

To calculate hydrophobicity surface properties, first we assign distinct hydrophobicity scales (BM, KD, EI and Wimley-White (WW)^54^), normalized between - 5 to 5, to each residue and their associated vertices in the mesh. Next, we recalculate the hydrophobicity values for each vertex by averaging the values of nearby surface mesh vertices within a distance cut-off of 10A. This local surface averaging process amplifies or attenuates a residue’s hydrophobicity score based on the hydrophobicity of its neighboring vertices, thereby resulting in the formation of hydrophobic or hydrophilic patches on the protein surface. The hydrophobicity surface scores for each scale (referred to as HPATCH) are summed for all the positive values within regions associated with each antibody domain (Fab, Fv, or CDR) (Figure 1C). This approach yields four distinct surface descriptors (one for each hydrophobicity scale) for each antibody domain. Note that while we generate descriptors on each antibody domain, here our primary focus is to examine the quality of surface representations specifically within the CDR region.This choice aligns with previous studies^1^ highlighting that the major differences across antibodies predominantly occur within the CDRs.

Figure 2 shows the correlations between HPATCH hydrophobicity scores across various scales, alongside the overlapping distributions for 674 clinical stage therapeutics (CST) and about 2000 human antibodies randomly selected from the Observed Antibody Space (OAS).^51^ It is evident that the different hydrophobicity scales are not perfectly correlated with one another for both CST and OAS. Notably, the EI and BM scales display the highest correlations with each other, followed by BM and WW. In contrast, the widely used KD scale demonstrates weak correlations with all three alternate scales. This observation contradicts the findings of Raybould et al.,^1^ who suggested strong correlations among these scales when using the PSH score. The observed discrepancy is likely due to the difference in how residues are classified as surface-exposed or buried between the PSH approach and our approach. While the PSH approach utilizes a fixed cutoff to define exposure, our approach considers the surface area associated with each atom in each residue. This allows our approach to account for the influence of large aromatic groups (Trp, Phe, Tyr) more prominently, leading to a stronger differentiation between hydrophobicity scales that rank aromatic residues as highly hydrophobic (BM, WW) compared to scales that do not (EI, KD). Moreover, our approach incorporates neighboring effects based on the proximity of residues on the surface, whereas the PSH approach relies on the closest heavy-atom Cartesian distance. This difference in considering neighboring effects may also contribute to the observed discrepancy between the two methods.

**Figure 2:**
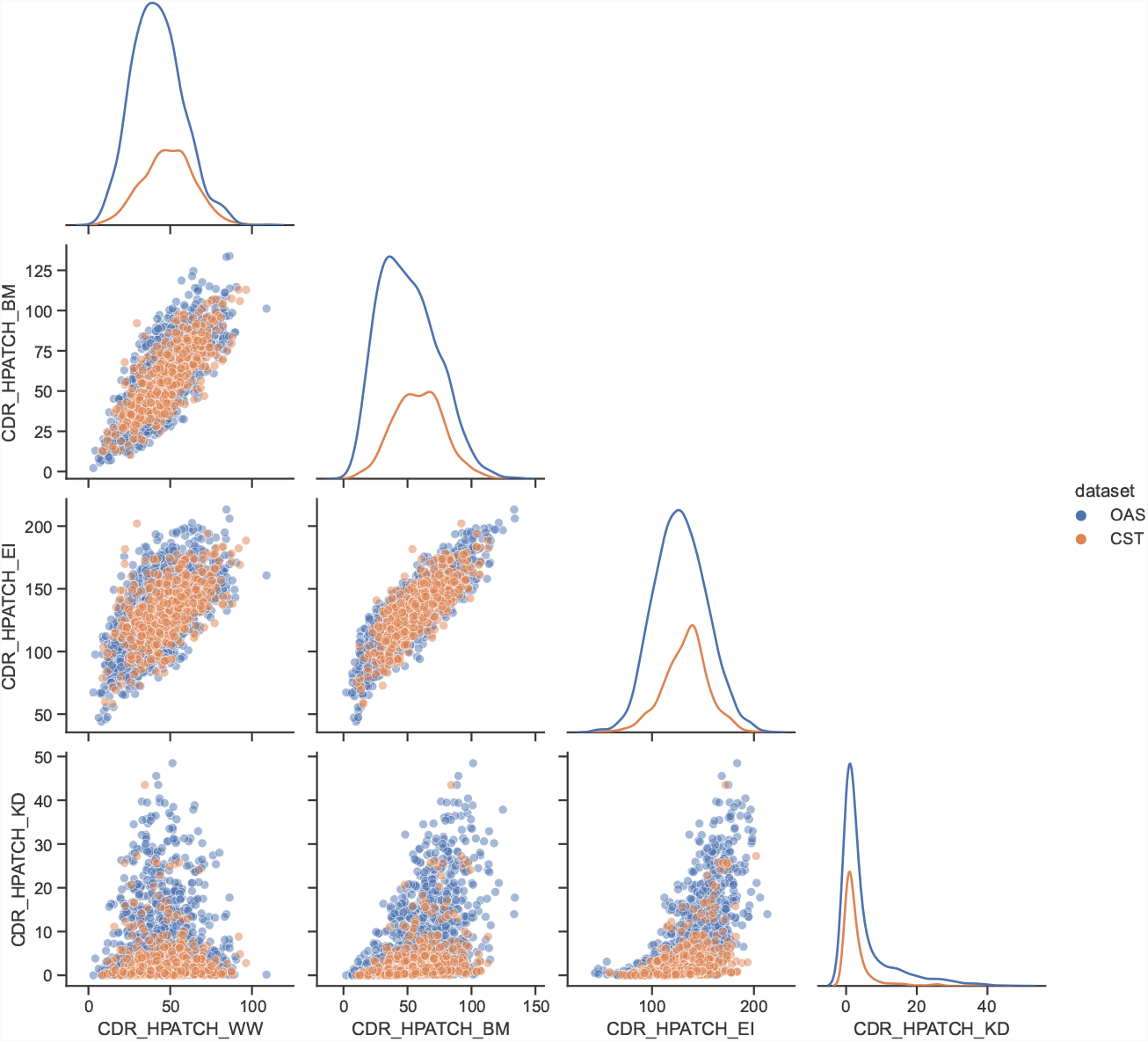
Matrix of scatter plots and distributions of the HPATCH hydrophobicity scores using four different scales (BM: Black & Mould, WW: Wimley-White, KD: Kyte-Doolittle, and EI: Eisenberg), for 674 clinical stage therapeutics (CST) and 2000 human antibodies randomly selected from the Observed Antibody Space (OAS). ^51^ The descriptor values are calculated based on the conformation of the initial structure model, AB2.

An illustration of the sensitivity to the choice of hydrophobicity scales is evident in Figure 2, where the different scales lead to inconsistent representations of clinical antibodies at higher HPATCH values on the CDR. Consistent with Raybould et al.,^1^ the KD scale implies that clinical antibodies are systematically less hydrophobic than human antibodies; the kernel density estimate (KDE) plot on the far right of figure 2 shows that the clinical antibody distribution is shifted slightly left (less hydrophobic) than the distribution of natural antibodies. However, the WW scale, which assigns greater weight to aromatic groups, suggests that clinical antibodies are more hydrophobic than human antibodies, as shown in the leftmost KDE plot in figure 2. This aligns with the broader engineering strategies employed to enhance potency by enriching binding sites with aromatic groups,^12,13^ potentially resulting in an overall increase in hydrophobicity for therapeutic antibodies.^55^ As seen in Figure 1, the largest hydrophobic patch for the WW and BM scales are on the binding site (variable domain) of an example fab structure, whereas the largest hydrophobic patch for the KD and EI scales are on the elbow of the fab.

These findings highlight the great impact of the chosen methodology and the key parameters on the outcomes and interpretations of the surface descriptors’ significance and predictive power. To gain a better understanding of the performance of the introduced electrostatics and hydrophobicity descriptors, we focus on evaluating their correlation with experimental data in the following sections, thereby assessing their direct relevance in developability assessment. Note that our objective is not to construct a multi-parameter model but rather to examine the direct correlations of individual descriptors.

### Viscosity

Subcutaneous delivery of antibodies necessitates high-concentration solutions to achieve the desired dosage.^57^ However, at these elevated concentrations, the self-association of antibodies can lead to unacceptably high viscosity levels, resulting in significant challenges during manufacturing and administration processes. Previous studies^58–61^ have proposed that the viscosity of high-concentration antibody solutions arises from molecular self-associations driven by both long- and short-range electrostatic interactions. Specifically, the *in silico* studies by Sharma et al,^29^ PfAbNet,^56^ and SCM^30^ has indicated that the negative charge on the variable domain of antibodies correlates with viscosity.

With this knowledge, we evaluated the performance of our negative electrostatic potential descriptor on the CDR, CDR_APBS_neg, calculated at pH 6.0, against two baseline methods: SCM^30^ and PfAbNet-viscosity.^56^ These methods were selected due to their reliance on Fv charge distribution and their specific design for viscosity prediction. SCM is a commonly used structure-based approach, while PfAbNet-viscosity is a 3D convolutional neural network (CNN) architecture trained to identify the key biophysical drivers of viscosity using the electrostatic potential surface of the antibody variable region as the sole input. To evaluate the performance, we utilized a dataset of 38 antibodies from Apgar et al.^62^ (PDGF38) which are mutants of the same parent molecule engineered to exhibit varying viscosity profiles;^62^ viscosity for this dataset was measured at pH 5.8 in the original paper. In line with the analysis in the PfAbNet-viscosity paper, we defined two classes using a threshold of 20 centipoise (cP): viscous (≥20 cP) and non-viscous (< 20 cP).

Figure 3 illustrates that CDR_APBS_neg at pH 6.0 outperforms the baseline methods SCM and PfAbNet-viscosity in terms of Pearson *R*^2^ (0.64, 0.60, 0.62, respectively), while PfAbNet has a higher Spearman *ρ* (0.78, 0.75, 0.80, respectively) (Figure 3C). All three models effectively differentiate between viscous and non-viscous antibodies in the dataset, with PfAbNet-viscosity performing slightly better (0.74, 0.80, 0.84, respectively). The comparable performance of our descriptor to a neural network explicitly trained for viscosity prediction suggests that a well-designed structural descriptor alone is capable of capturing this type of physical behavior effectively.

**Figure 3:**
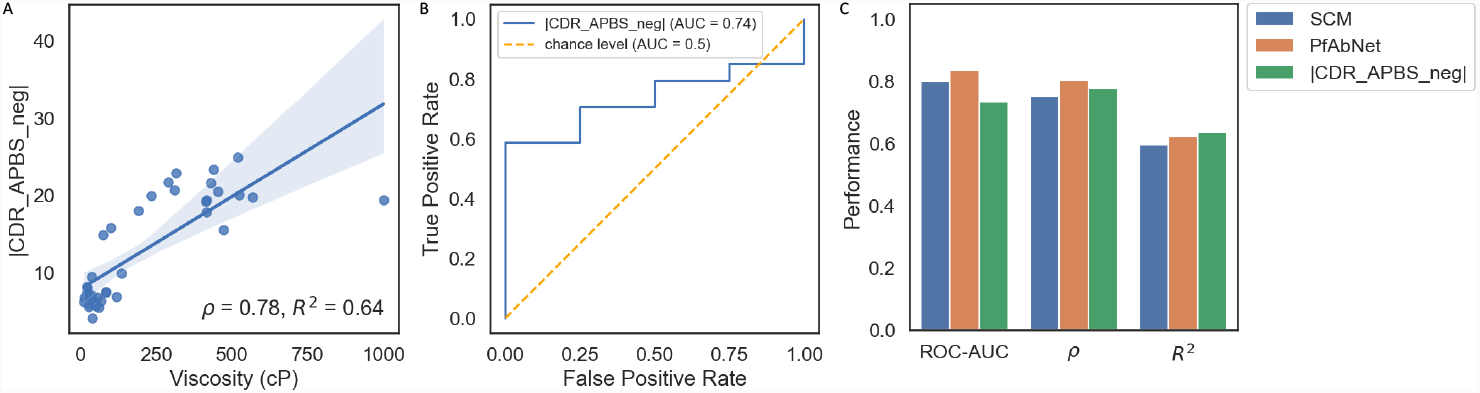
(A) Viscosity data for 38 antibodies from Apgar et al. (PDGF38), which are variants of the same clone with modulating viscosity, ^56^ plotted against the absolute value of CDR_APBS_neg at pH 6.0, showing a Spearman *ρ* of 0.78 and Pearson *R*^2^ of 0.64. (B) A Reciever-Operator-Characteristic (ROC) curve demonstrating an area-under-curve (AUC) value of 0.74 when predicting a binary viscosity label calculated using a threshold of 20 cP. (C) Comparisons of ROC-AUC, Spearman *ρ*, and Pearson *R*^2^ for the absolute value of CDR_APBS_neg at pH 6.0, SCM, and PfAbNet against the same viscosity dataset (PDGF38), which demonstrates higher Pearson *R*^2^ correlation with our descriptor than either, a marginally lower Spearman *ρ* than PfAbNet, and a lower ROC-AUC than both SCM and PfAbNet.

### PK Clearance, Polyspecificity and Heparin binding

Pharmacokinetic (PK) properties of antibody drugs impact efficacy, dose, and dose intervals.^66,67^ In particular, fast PK clearance, or short drug half-life, can lead to issues with patient compliance and clinical benefit. Several studies have indicated that non-Fc related differences in PK clearance can be the result of off-target binding,^66–68^ or polyspecificity, which has been linked to high hydrophobicity and extreme charge distributions on the variable domain.^9,29,67^ Sharma et al.^29^ used two sequence-based descriptors (the net charge of the Fv at pH 5.5 and the sum of the hydrophobic index along several CDRs), to determine nonspecific binding and predict fast clearance. More recently, Grinshpun et al.^63^ curated a comprehensive clinical PK dataset with calculated estimates of linear clearance for 64 antibodies, and demonstrated that using structure-based descriptors, such as structure-based pI (pI_3D) and patch_cdr_hyd from MOE, can improve the discrimination of fast clearing antibodies.

To evaluate the applicability of our electrostatic and hydrophobicity descriptors in this context, we evaluate the performance of the sum of positive electrostatic potential on the surface of the CDR region (CDR_APBS_pos at pH 7.4) and the hydrophobicity calculated using HPATCH and the Wimley-White scale (CDR_HPATCH_WW) in discriminating fast clearing antibodies in the same set of 64 clinical stage antibodies mentioned above (Figure 4A-D). To determine the relevant thresholds that correspond to non-extreme ranges for these two parameter space, we performed a two-dimensional kernel density estimation on a dataset comprising 674 clinical antibodies (Figure 4B). The dark contour values in the most populated region represent approximately 68% of the data and fall within the range of 7 to 36.5 for CDR_APBS_pos, and 0 to 80 for CDR_HPATCH_WW (Figure 4C).

**Figure 4:**
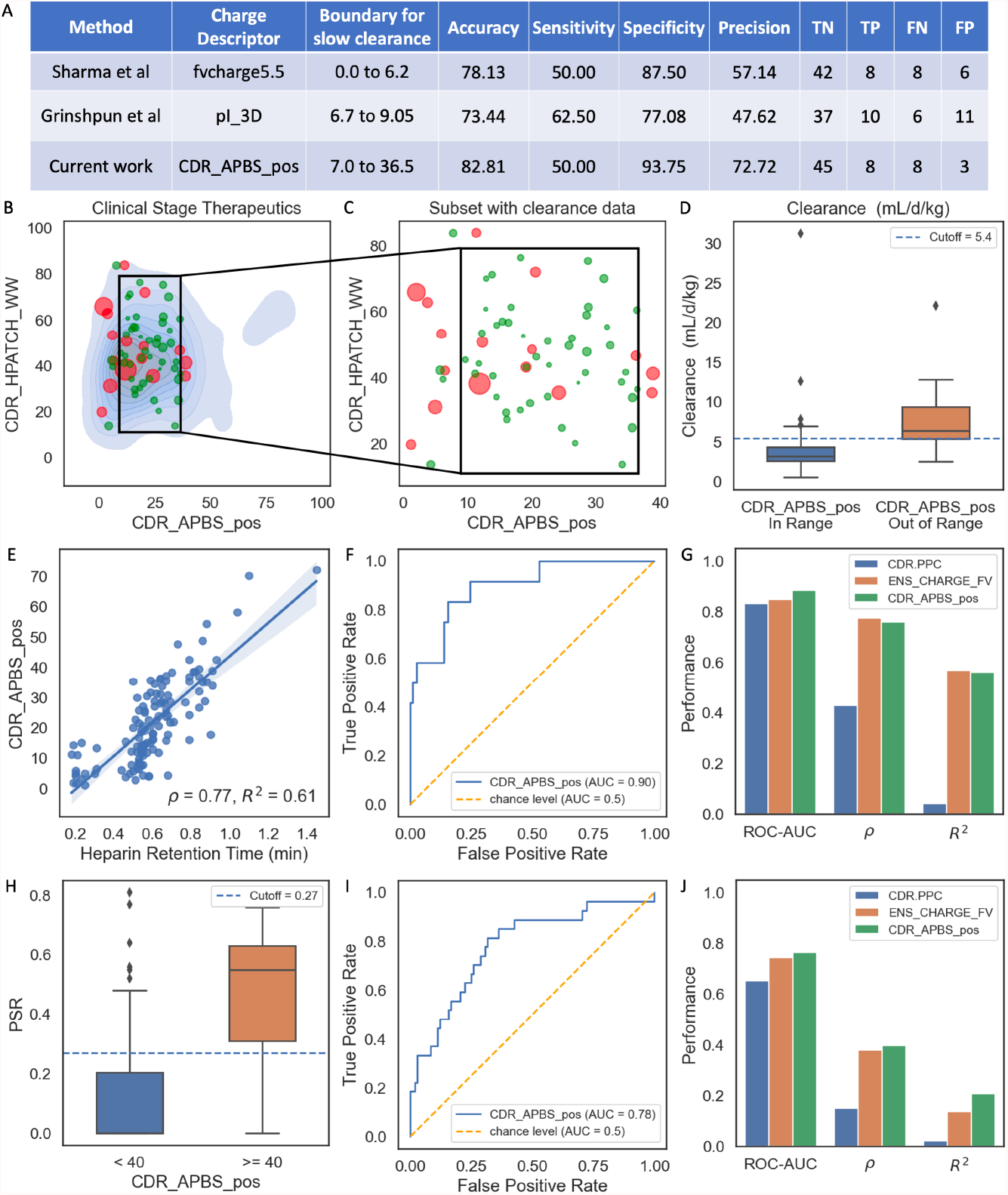
A) Table comparing CDR_APBS_pos at pH 7.4 with Fv Charge at pH 5.5^29^ and pI_3D, ^63^ using confusion metrics against PK-clearance data. ^63^ B) The distribution of clearance data on a CDR_APBS_pos at pH 7.4 vs CDR_HPATCH_WW scatterplot with the blue KDE in the background showing the 2D distribution for all CSTs; dot-size represents relative clearance rate (mL/d/kg) and red represents fast clearance. C) Partition shows high clearance molecules falling primarily outside of 2D box. D) A boxplot showing clearance distributions for molecules in- and out-of-range of CDR_APBS_pos thresholds. Performance of CDR_APBS_pos at pH 7.4 in predicting Heparin RT ^64^ (E-G), and PSR data ^65^ (H-J) relative to TAP’s ^1^ CDR.PPC and MOE’s ^31^ ENS_CHARGE_FV.

In line with the findings of Grinshpun et al.,^63^ we observed that hydrophobicity does not make a significant contribution to differentiating antibodies with fast and slow clearance, at least in this dataset (Figure 4C), unlike what was reported by Sharma et al.^29^ One possible explanation for this inconsistency could be due to elimination or optimization of extremely hydrophobic attributes in these clinical molecules, prior to progressing to the clinic. Given that the electrostatic-based descriptors play a more prominent role in this discrimination, we focused our benchmarking solely on comparing the predictive power of different charge-based descriptors: CDR_APBS_pos at pH 7.4, fvcharge5.5 and pI_3D (Figure 4A). For the reference descriptors we used the thresholds established by Grinshpun et al.^63^ Figure 4A demonstrates that by utilizing charge-based descriptors with listed thresholds as binary predictions, CDR_APBS_pos improves the discrimination of antibodies with fast clearance. It correctly excludes 8 out of 16 antibodies with fast clearance (≥5.4 mL/day/kg^63^) and accurately predicts 45 out of 48 antibodies with slow clearance (<5.4 mL/day/kg^63^). CDR_APBS_pos predicts binary clearance with 82.81% accuracy, an increase relative to the other two baseline descriptors (78.13 and 73.44 for fvcharge5.5 and pI_3D, respectively).

Poly-specificity reagent (PSR) and heparin chromatography are two early-stage developability assays that have been associated with poor PK properties of antibodies, and both have been shown to correlate with the electrostatics, particularly positive charge patches on the Fv.^64,65^ Here we use CDR_APBS_pos at pH 7.4 and benchmark it against two previously studied electrostatics-based descriptors, including MOE’s ens_charge_Fv^69^ and Raybould et al.’s Therapeutic Antibody Profiler (TAP) CDR Positive Patch Charge (CDR.PPC),^1^ in predicting both PSR and Heparin binding using the *in silico* and *in vitro* data curated by Jain et al. (2023).^64,65^

With respect to the PSR assay data, we observe improvement in ROC-AUC, Spearman rho, and Pearson *R*^2^ (0.76, 0.4, 0.15 respectively) over both MOE’s ens_charge_Fv (0.74, 0.38, 0.14 respectively) and TAP’s CDR.PPC (0.65, 0.16, 0.03 respectively) (Fig 4H-J). For heparin chromatography data, our CDR_APBS_pos had a higher ROC-AUC score than both MOE’s ens_charge_Fv and TAP’s CDR.PPC (0.89, 0.85, 0.83 respectively) (Fig 4F-G). For both *ρ* and *R*^2^ (Fig 4E-G), MOE’s ens_charge_Fv outperforms our CDR_APBS_pos by a small margin, while both significantly outperform TAP’s CDR.PPC. The improvement across the board, in the case of PSR, and the similarity in performance with MOE in the case of heparin chromatography, highlights the ability of our descriptors to capture polyspecificity.

### Aggregation and HIC retention time

Aggregation presents a significant obstacle to the advancement and development of therapeutic monoclonal antibodies (mAbs). The aggregation propensity of mAbs is often associated with the hydrophobicity of their CDRs (SAP,^32^ PSH,^1^ CamSol,^70^ Aggscore^33^). In order to assess the predictive power of our hydrophobic descriptors (specifically HPATCH applied to the CDR using different hydrophobicity scales), we conducted a correlation analysis with two *in vitro* screening assays.

First, we analyzed a set of 13 antibodies (internal Genentech molecules) with size exclusion chromatography (SEC) data capturing % monomer loss, a surrogate metric for aggregation, see Methods and Figure 5A-C. Among these antibodies, six exhibited high % monomer loss values, indicating aggregation, while the remaining seven behaved well (.2 is an internal threshold used for this assay). When partitioning the groups by a hand-tuned CDR_HPATCH_WW threshold (43), we see good separation with respect to the aforementioned % monomer loss cutoff. As shown in Figure 5C, CDR_HPATCH_WW and BM outperformed the other metrics in terms of correlation. Interestingly, CDR_HYD from MOE, which employs a LogP-based hydrophobicity scale, displayed a negative correlation. This observation further supports the notion that LogP-based hydrophobic patches might not be directly relevant in the context of protein-protein interactions that promote aggregation.

**Figure 5:**
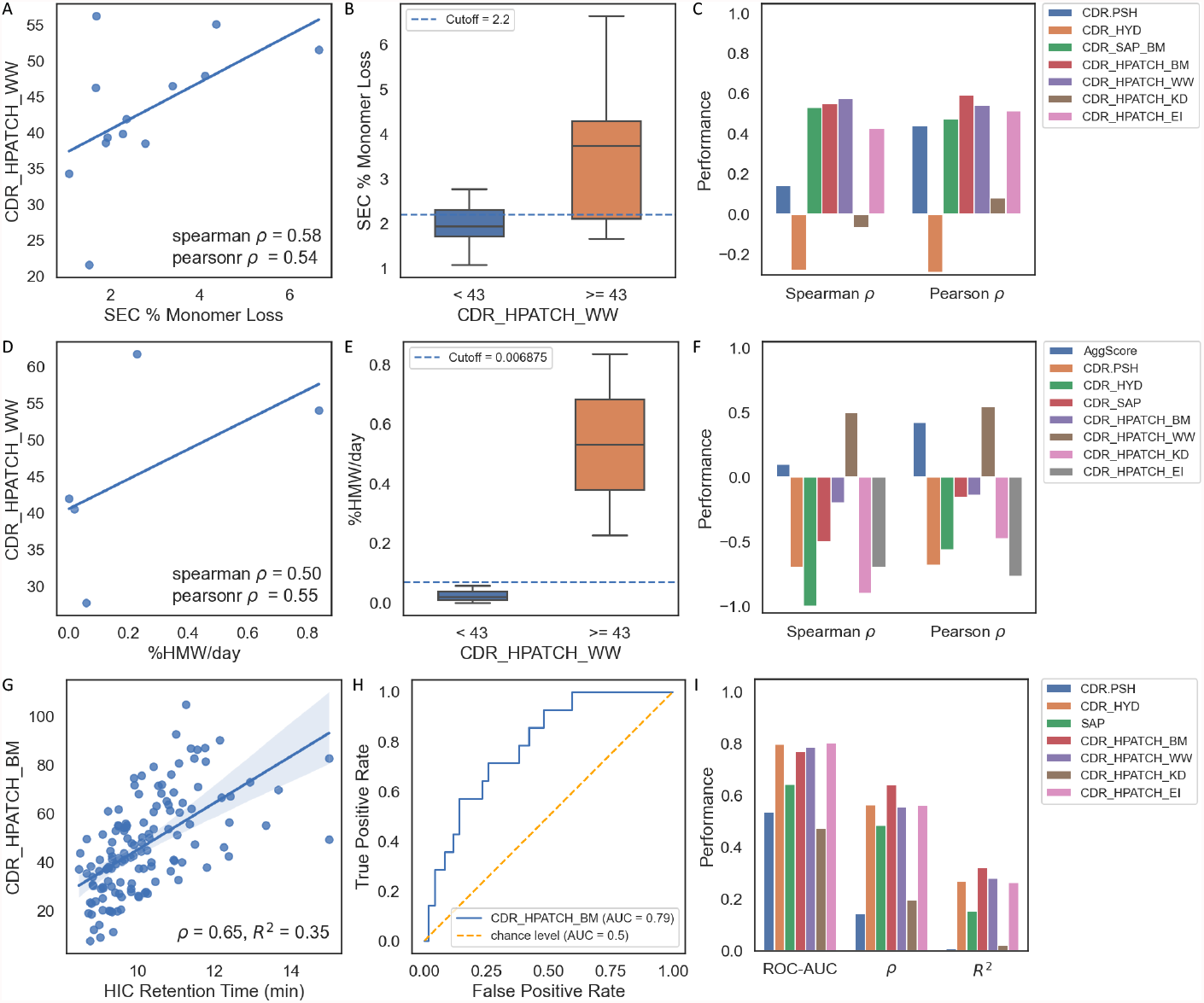
Performance of CDR_HPATCH_WW against RGNE-generated SEC data (% Monomer Loss) as: a scatterplot (A); a boxplot showing % Monomer Loss distributions for molecules above and below a CDR_HPATCH_WW threshold of 43 (B); and comparison of Spearman and Pearson *ρ* of % Monomer Loss with TAP’s CDR.PSH, ^1^ MOE’s CDR_HYD, ^31^ an inhouse implementation of SAP, ^32^ and MolDesk CDR_HPATCH with BM, WW, KD, and EI scales (C). Same set of analysis as A-C, but for SEC %HMW/day as generated for Highland Games 2020^34^ (D-F), with an added comparison of AggScore. ^33^ Performance of CDR_HPATCH_BM with HIC RT as a scatterplot (G), ROC curve (H), and performance of CDR_HPATCH with all hydrophobicity scales against *in silico* descriptors listed previously (I).

Next, we examined the correlation between HPATCH scores, CDR_HYD,^31^ CDR_PSH,^1^CDR_SAP,^32^ AggScore,^33^ and the percentage of high molecular weight species generated per day (%HMW/day) for a set of five antibodies from an antibody property prediction competition, Highland Games 2020^34^ (5D-F). Among these antibodies, two exceeded the threshold, 0.006875 (defined in the Highland Games paper^34^) for %HMW/day, while the remaining three exhibited minimal aggregation. We see perfect separation of groups with respect to the aforementioned %HMW/day threshold when partitioning the data by the same CDR_HPATCH_WW threshold (43) as used for SEC %monomer loss. Surprisingly, all descriptors, except AggScore and CDR_HPATCH_WW, displayed a negative correlation with %HMW/day (Figure 5F), falsely indicating that higher hydrophobicity was associated with lower aggregation rates.^59^ Notably, WW was the only hydrophobicity scale resulting in a positive correlation, suggesting the involvement of aromatic residues in modulating protein-protein interactions leading to the formation of high molecular weight aggregates.

Hydrophobic interaction chromatography (HIC) is an analytical method used to separate molecular variants of proteins according to surface hydrophobicity which can be used to bolster the developability profile of therapeutic candidates.^71,72^ Using the HIC data generated for 137 mAbs by Jain et al. in 2017,^73^ we compare the predictive performance of our *in silico* HPATCH descriptors against TAP’s CDR.PSH, MOE’s CDR_HYD, and SAP (Figure 5G-I).

We observe higher Spearman *ρ* and Pearson *R*^2^ with our CDR_HPATCH_BM descriptor than CDR.PSH, CDR.HYD, and SAP (Figure 5I). CDR_HYD, which is tuned specifically for HIC, marginally outperforms our descriptors when compared using ROC-AUC (Figure 5I)).

### Optimal choice of interior dielectric constant

The dielectric constant for bulk water in equilibrated systems is typically agreed to be 78.4.^74^ However, determining the optimal dielectric constant for the interior regions of proteins is a debated topic, with suggested values ranging from as low as *ε* = 1 or 2 to as high as *ε* = 40.^74,75^ To determine the optimal interior dielectric for representing electrostatics of antibody variable domains, we examined the correlation of the CDR_APBS_pos descriptor to Heparin and PSR data (Figure 6A). We screened a range of interior dielectric constants from 1 to 20, with a fixed exterior dielectric of 78.4. The results showed that the choice of interior dielectric constant significantly affected the correlation between the electrostatic positive patch on the CDR and Heparin/PSR data. Higher dielectric constants improved the Pearson correlations by as much as 20%, with improvement plateauing at an interior dielectric constant of 16 (further increases did not notably improve the correlation). Figure 6B also shows that the visualization of the electrostatic potential on the surface of Omalizumab (as an example) becomes more refined and detailed with higher dielectric values. These findings stress the importance of appropriate dielectric constants for accurate interpretation of molecular electrostatic properties of antibodies. Based on the benchmark analysis here, we used an interior dielectric of 16 for all our analysis.

**Figure 6:**
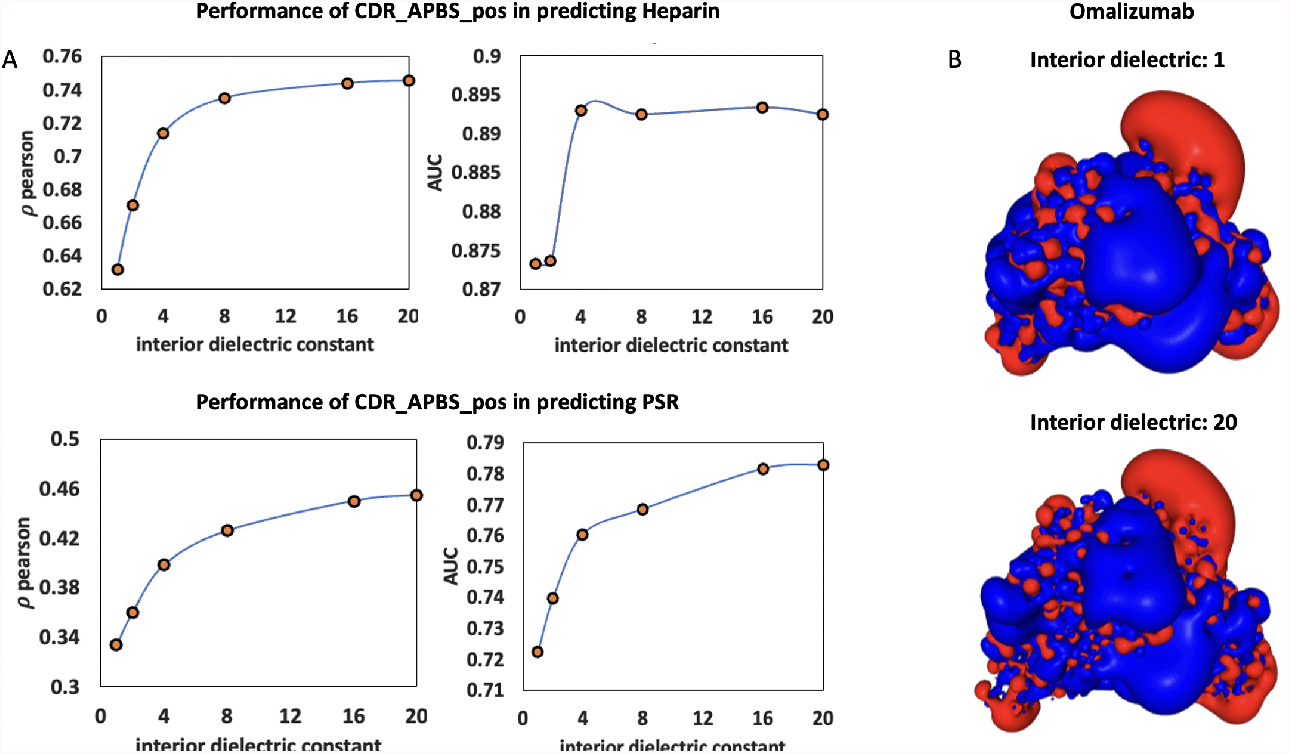
A) Impact of Interior Dielectric Constant on Correlation of Electrostatic Positive Patch on the CDR with Heparin and PSR Data. B) Visualization of Electrostatic Potential on the Surface of Omalizumab at Extreme Interior Dielectric Values (0 and 20). The descriptor values are calculated based on the conformation of the initial structure model (AB2) at pH 7.4.

### Sensitivity to the antibody structure model

The accuracy of structure and surface-based descriptors and properties are heavily dependent on the quality of antibody structure model^43,47^ Several methods have been developed to improve antibody structure predictions, including deep learning approaches such as ImmuneBuilder Antibody Builder 2 (AB2),^44^ DeepAb,^45^ Equifold,^46^ and homology modeling methods such as MOE.^31^ While these models have primarily been evaluated for predicting X-ray crystal structures, their impact on the molecular surface properties relevant to antibody developability, as well as their comparative accuracy, remain unclear.

To address this, here we conducted a benchmark analysis using four different initial structure models (AB2,^44^ DeepAb,^45^ Equifold^46^ and MOE^31^), and examined their impact on the correlation between our descriptors and empirical developability data. Figure 7A-B shows the Pearson *R*^2^ and ROC-AUC values obtained for predicting various *in vitro* assays using corresponding molecular descriptors. Overall, we observed similar performance across the different *in vitro* datasets for the tested structure models. While no model consistently outperformed the others across all datasets, AB2 demonstrated greater accuracy in viscosity prediction, MOE performed well in predicting PSR, and DeepAb showed slightly better correlation with HIC. Therefore, we conclude that without conformational sampling, all tested single-static structure models yield comparable accuracies.

**Figure 7:**
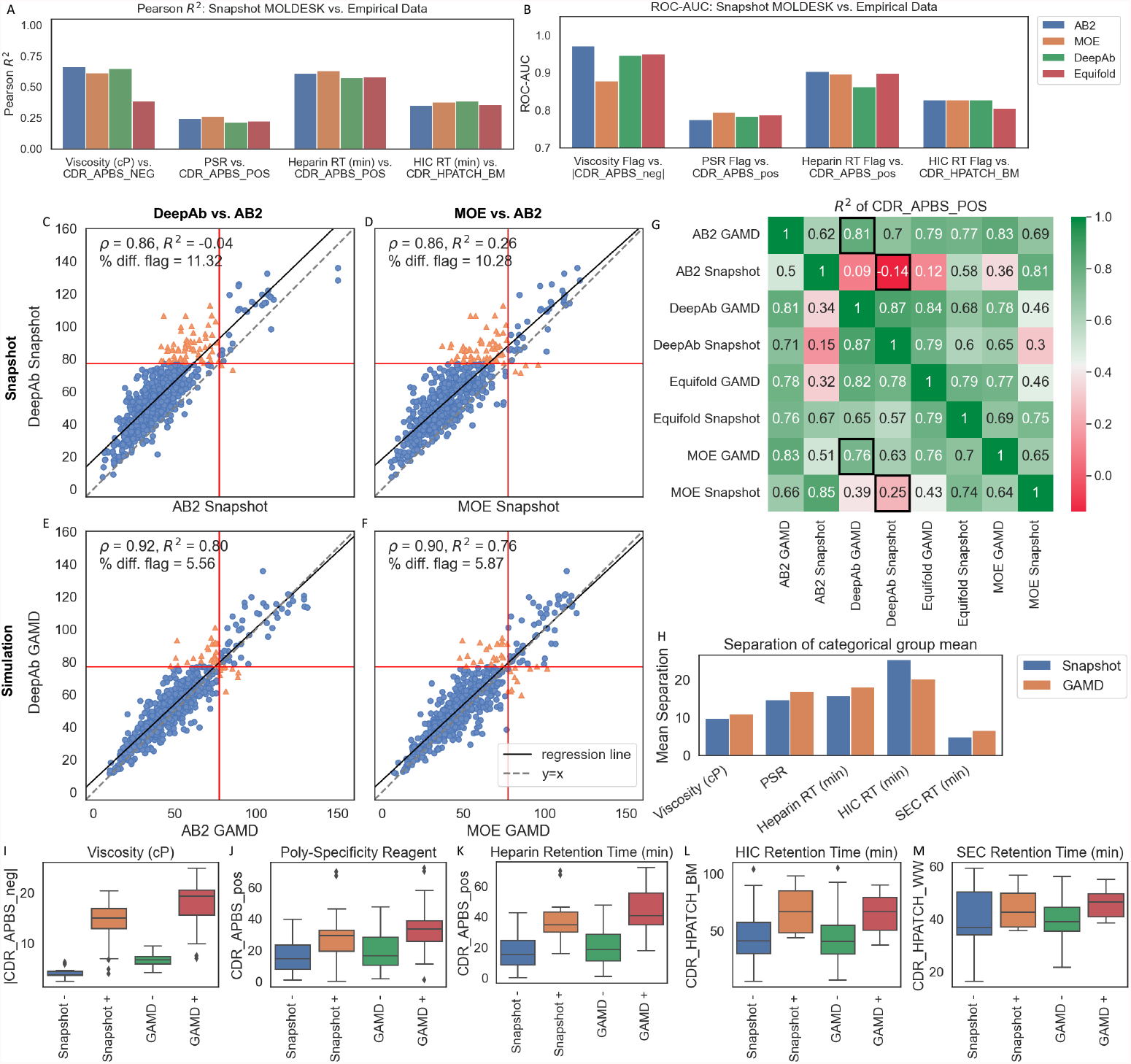
Pearson *R*^2^ correlations (A) and ROC-AUC scores (B) of AB2, MOE, DeepAb, and Equifold snapshot Fv structures against viscosity, PSR, heparin retention time, and HIC retention time. Figures C-F show correlation plots of CDR_APBS_pos with 10 percentile cutoffs (red vertical and horizontal lines) calculated in advance. Blue circles indicate agreement in risk category for the x and y sets and orange triangles indicate a mismatch in risk category, with Spearman *ρ*, correlation of determination, *R*^2^, and the % of mismatched flags according to the cutoff. G) A heatmap showing the coefficient of determination, *R*^2^, for CDR_APBS_pos across AB2, MOE, DeepAb, and Equifold sets for both single-static initial structure and ensemble-averaged values (5ns of GaMD simulations). I-M) After partitioning *in vitro* datasets into high (+) and low (-) risk categories according to thresholds defined in their original publications, we compare boxplot distributions of the corresponding MolDesk descriptors generated using single-conformation (snapshot) initial structures and GaMD sampling. H) a barplot shows the difference in mean values between the high and low risk distributions for both sampling modes: all except for HIC RT show greater separation in distributions after applying conformational sampling.

To gain further insights into the distinctions between different initial structure models, we calculated the coefficient of determination, *R*^2^ of CDR_APBS_pos values across a panel of 557 clinical antibodies using each of the four models as initial structure models. All columns and rows in Figure 7G with “Snap-shot” labels denote the initial structure models with no sampling. Surprisingly, the cross-correlation analysis suggested that different models could lead to distinct descriptors with systematic shifts in the value distribution, resulting in *R*^2^ correlations as low as - 0.14 between AB2 and DeepAb. The best correlation between static structure models was observed between AB2 and MOE, while the weakest correlation was found between AB2 and DeepAb. This finding is intriguing because MOE is based on fragment-based homology modeling, whereas both DeepAb and AB2 rely on deep learning. One expects a better agreement between DeepAb and AB2, especially considering that the structures used in training both models were from the Structural Antibody Database (SAbDab).^44,45^ These results show that although these weak correlations did not manifest in the predictive performance for the aforementioned *in vitro* datasets, they could lead to different risk flags when using a specific cutoff threshold (see section below on risk flags). For example, Figure 7C shows that for CDR_APBS_pos calculated from initial structure model outputs, DeepAb-based analysis results in a systematically higher number of risk flags (above the horizontal red line) than AB2 (to the right of the vertical red line). This highlights the sensitivity of developability metrics, particularly risk flags, to the choice of initial structure models when a single-static structure is used.

### Impact of conformational sampling

Next, we sought to examine whether the substantial disagreement observed between the initial structure models could be rectified through conformational sampling. To investigate this, we generated conformational ensembles by performing 5 ns of GaMD simulations on each set of 557 Fvs generated by the four structure models. Our analysis revealed that when the values for the CDR_APBS_pos descriptor were averaged over a conformational ensemble, improved correlations were observed among different structure models compared to those calculated using single-static structures (Figure 7G). Additionally, Figures 2E-F demonstrated an enhanced correlation between the classification of the CDR_APBS_pos binary risk flags, using a cutoff threshold of 10 percentile (see section below), when the values were averaged over a conformational ensemble. Notably, the *R*^2^ correlation between Ab2 and DeepAb improved from - 0.14 to 0.81 (Figures 7C and E), while the *R*^2^ correlation between DeepAb and MOE increased from 0.25 to 0.76 (Figures 7D and F) following conformational sampling. In both cases, the percentage of mismatched flags (% diff. flag) was reduced by half after conformational sampling, from 11.32% to 5.56% for DeepAb vs. AB2 (Figures 7C and E) and from 10.28% to 5.87% for DeepAb vs. MOE (Figures 7D and F). These findings suggest that with sufficient conformational sampling, the accuracy of surface descriptors from different structure models converge.

To further explore the direct impact of conformational sampling on the accuracy of predictions for *in vitro* assays, we conducted additional analyses using AB2 as the initial model. We focused on evaluating the categorical predictions of several different assays (Figures 7H-M). Our findings indicate that conformational sampling consistently enhances the separation of categorical predictions for all the different assays, with the exception of HIC. This observation aligns with the results reported by Waibl et al.,^41^ suggesting no systematic improvement in predicting HIC due to sampling. We speculate that this decrease could be attributed to either insufficient sampling or an overall lower quality of the HIC dataset. Nevertheless, our data supports the conclusion that conformational sampling systematically improves the accuracy of predictions for various developability attributes and *in vitro* assays.

Based on this analysis, we propose that all the different structure models studied here perform equally well in predicting descriptors relevant to developability prediction, as long as conformational sampling is carried out and there are no inherent structural issues that cannot be fixed through conformational sampling (see Ref.^47^ for a detailed discussion of such structural issues). Furthermore, after assessing the quality of structures using TopModel^47^ (data not shown), we recommend utilizing AB2 as the preferred model since it consistently generates fewer faulty structures and is fast and freely available.

### Developability risk flags and ability to predict progression in the clinic

In addition to calculating molecular descriptors and exploring correlations with *in vitro* data, *in silico* studies have proposed strategies to flag monoclonal antibodies (mAbs) as high or low risk for development based on the number of violations of these descriptors.^1,69,76^ Notably, Raybould et al. introduced five computational guidelines known as the Therapeutic Antibody Profiler (TAP) to evaluate the developability of mAbs.^1^ These guidelines involve analyzing the distribution of molecular descriptors in a reference set comprising clinically approved mAbs and representative mAbs from the human repertoire. By identifying mAbs that fall at the extremes or tails of these descriptor distributions, potential risks can be flagged.

Furthermore, several studies have demonstrated a correlation between the number of developability risk flags, whether derived from *in vitro* or *in silico* assessments, and the various stages of clinical progression. For example, Jain et al. conducted a study^65^ where they observed a decrease in the number of risk flags, as indicated by *in vitro* assays such as PSR and HIC, as a drug progressed through clinical trials, ultimately leading to its approval. While the primary determinants of clinical success lie in the biological function and efficacy of the antibody, the association between the number of developability risk flags and the progression in clinical stages is intriguing, as thoroughly discussed in the work by Jain et al.^65^

To evaluate the discriminatory ability of the descriptors introduced in this study for clinical antibodies that either progressed or regressed in clinical stages, we selected six descriptors from our set based on their predictive power for various experimental liabilities as shown in previous sections (Figure 8B). These descriptors include CDR_APBS_neg, a significant factor influencing viscosity and colloidal stability (Figure 3), CDR_APBS_pos, a major driver of PK clearance and polyspecificity (Figure 4), CDR_HPATCH_WW, which plays a key role in SEC, HMW, and HIC (Figure 5), CDR_HPATCH_BM which correlated the best with HIC, as well as total CDR length and APBS charge asymmetry (APBS_CAP) as two additional descriptors to align with the type of descriptors considered in the TAP metrics.^1^

**Figure 8:**
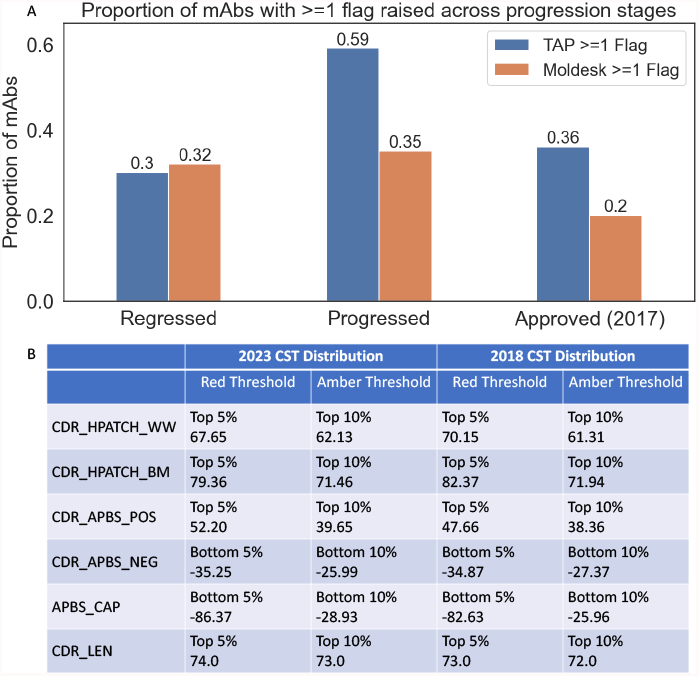
(A) Proportion of mAbs with ≥ 1 risk flag raised using MolDesk and TAP’s flagging systems for molecules which have regressed (failed for any reason or moved down in clinical trial status since 2017), progressed (approved or moved into next clinical phase since 2017), and those which were approved in 2017. Flagging thresholds were determined using a subset of 238 CSTs from 2018^1,65^ for both MolDesk and TAP, shown in (B) alongside the MolDesk thresholds computed from 629 clinical stage molecules in 2023. ^77^

To identify extreme ranges and risk flags, we utilized distributions of corresponding descriptors for a subset of clinical antibodies from 2018 (as used in the TAP paper,^1^ refer to Adimab 2023^65^) and defined the flags using the 5% tails of these distributions for both TAP and MolDesk metrics. We specifically chose clinical antibodies from 2018 to avoid any bias in our thresholds resulting from the inclusion of more recent molecules, similar to the study by Jain et al.^65^ Subsequently, we assessed the accuracy of the TAP five metrics and our descriptors in predicting clinical progression stages. For this analysis, we focused on 73 mAbs categorized as either Progressed or Regressed in the clinic, as described by Jain et al.^65^ (Figure 8A illustrates the fraction of mAbs with violations for the egressed, Progressed, and Approved 2017 sets of mAbs). For each class, we further classified the mAbs into different categories based on the presence or absence of flags using both TAP and MolDesk metrics.

Figure 8 illustrates the distribution of these classifications. Interestingly, we observed that both metrics performed almost similarly in identifying regressed molecules, with MolDesk showing slightly higher proportion of mAbs with at least one risk flag. However, when it came to predicting progression in clinical antibodies, MolDesk significantly outperformed TAP. Specifically, while TAP flags assigned 1 or more flags to 59% of of progressed molecules, MolDesk flags identified 1 or more flags in only 35% of these molecules. Similarly, while TAP assigned flags to 36% of approved molecules, MolDesk flags assigned 1 or more flags to 20% of them. The smaller number of risk flags by MolDesk assigned to both progressed and approved set, with a similar performance on the regressed molecules, highlights the superior predictive capability of the MolDesk metric in identifying mAbs that are more likely to progress successfully through clinical stages, emphasizing its potential as a valuable tool for assessing developability.

## Discussion and concluding remarks

In this work, we introduced MolDesk, a set of molecular surface representations specifically designed to predict antibody developability characteristics. We evaluated the performance of these representations by comparing their correlations with a diverse range of *in vivo* and *in vitro* data. The MolDesk surface descriptors were developed to capture the essential biophysical properties of antibody variable domains, including electrostatics and hydrophobicity features. To represent electrostatics, we employed a direct mapping approach, where the continuous electrostatic potential from a cubic grid was projected onto a surface mesh and subsequently to the residues and specific antibody domains (CDR, Fv, etc). This enables us to calculate ensemble-averaged surface electrostatic potential values per residue and per domain, providing a more comprehensive and dynamical representation of the antibody’s electrostatic properties compared to the traditional qualitative approach of visualizing electrostatic potentials using APBS at a single conformation. Additionally, by carefully selecting protocol parameters, such as the interior dielectric constant, we observed improvements in agreement with empirical data that has been previously correlated with electrostatic potential, namely PK Clearance, Heparin and PSR.

An interesting finding was that the negative electrostatic surface potential on the CDR (CDR_APBS_neg) performed comparably to a neural network model specifically trained on electrostatic surface potential for high-concentration viscosity prediction. This suggests that a finely-tuned structural descriptor alone is sufficient for capturing this particular physical behavior. In this context, the quality of input structure-based features to machine learning algorithms becomes equally important to the model training process itself. Furthermore, the high-quality structural features calculated in our approach are valuable for developability analysis due to their interpretability and independence from the quality of training data. This is particularly relevant as developability-related assays often involve small and noisy datasets.

Additionally, our hydrophobicity-based descriptors (HPATCH) offer several improvements over previous approaches. Firstly, we incorporated the average hydrophobicity values of neighboring surface mesh vertices to account for the clustering effect of similarly-typed residues. Secondly, instead of relying on a pre-selected hydrophobicity scale, we tested multiple commonly used scales and identified the optimal choices based on their correlations with hydrophobicity-driven developability assays. Our results supported the superior predictive power of the BM and WW scales in predicting experimental assay results relevant to aggregation (SEC %HMW/day, % monomer loss) and HIC RT.

We also investigated the impact of the initial structure models’ quality on the accuracy of our surface descriptors. Our findings revealed that when using static initial structures, direction correlations between MolDesk descriptors based on different model structures (AB2, DeepAb, Equifold, and MOE) and empirical data were comparable. However, the descriptors derived from static structures generated using different initial models did not correlate well with each other. When ensemble-averaged values were used instead of single-static structure values, we observed improved correlations between descriptors derived from different initial structures, which translated to improved discriminatory power in predicting empirical data as well. These results highlight the inherent insufficiency of single-static structures in accurately capturing antibody properties. Regardless of the quality of the input structure, relying solely on single-static structures can yield inconsistent and noisy outcomes. Employing conformational ensembles in solution is strongly recommended as a more reliable approach. It would be valuable to investigate the sensitivity of our results to different conformational sampling methods, including variations in simulation length or type. Additionally, future studies should aim to benchmark the accuracy of various initial antibody structure models through rigorous and detailed conformational sampling techniques.

Expanding on the well-established correlations between MolDesk descriptors and experimental developability data, we proposed six developability risk flags for profiling antibodies and guiding candidate selection through developability assessment. Notably, our descriptors demonstrated superior predictability of clinical progression compared to the established TAP rules. This finding highlights the practical applicability of our method, especially in the advanced stages of candidate selection, where prioritizing accuracy outweighs considerations of throughput and computational speed.

Future work can incorporate multi-parameter models that combine different descriptors proposed here to enhance correlations with specific experimental data. Moreover, we believe that our framework can be extended to define novel patch definitions by integrating and combining values on the vertex nodes using different representations. This approach holds the potential to unlock additional insights and refine our understanding of antibody surface properties in the context of developability assessment. Finally, while our study primarily focused on the application of these descriptors in the context of antibody developability, their utility could extend to improving the accuracy of surface-based predictive models for studying molecular recognition and protein-protein interactions.

## Methods

### Antibody structure modeling

By default, we utilized the Immunebuilder^44^ Antibody Builder 2 (AB2) structure prediction tool to generate Fv domain antibody structures. To evaluate the sensitivity to the choice of model structures, we also modeled Fv domain structures using: MOE^31^’s homology modeling tool; Equifold,^46^ a deep learning driven structure prediction tool; and DeepAb,^45^ a deep learning structure prediction method.

### Antibody preparation

The structure models were energy minimized using SANDER available in Amber 2020.^78^ PDB2PQR^79^ (v. 1.8) was used with the AMBER force field^80^ to assign partial atomic charges whereby ionization states were determined using PROPKA^81,82^ at pH 6.0 or 7.4 depending on the empirical assay studied. The resulting prepared structures were saved in PQR format.

### Molecular Dynamics Protocol

The Fv structures were parameterized with FF14SB force field,^83^ and were solvated in a cubic box of TIP3P explicit water model with a minimum 10Å distance from the edge of the box. The system charge was neutralized with Na+ and Cl-counter ions. Hydrogen Mass Repartitioning^84^ was performed on the solute atoms to enable a simulation time step of 4fs.

The GPU implementation of Amber 2020 MD software package^78^ with the SPFP precision model^85^ was used for all the MD simulations. First, the structure was relaxed with 2000 steps of conjugate-gradient energy minimization, using harmonic restraining potential with the force constant of 10 kcal/mol/Å^2^ to restrain the solute to the initial structure. Then the pressure was maintained at 1 atm and the thermostat temperature increased to 300K over the course of 200 ps keeping the Harmonic positional restraints of strength 10 1 kcal/mol/Å^2^ on the solute structure. The system was then equilibrated for 1 ns with a restraint force constant of 1 kcal/mol/Å^2^. All restraints were removed for the production stage. The simulation time step was 4 fs. A 9Å cutoff radius was used for range-limited interactions, with Particle Mesh Ewald^86^ electrostatics for long-range interactions. The production simulation was carried out using NPT conditions. Langevin dynamics^87^ was used to maintain the temperature at 300K with a collision frequency of *γ* = 1 ps^*−*1^.

After the 1 ns NPT equilibration, a 5 ns GaMD^52^ simulation module in AMBER 2020 was performed. All GaMD simulations were run at the dual-boost level, with the reference energy set to the lower bound. System potential energies were averaged and their standard deviation calculated every 50,000 steps (100 ps). The upper limit of the boost potential’s standard deviation, *σ* 0, was set to 6.0 kcal/mol for both the dihedral and total potential energetic terms. Simulation frames were saved every 20 ps for analysis.

### Electrostatic surface properties

To analyze the electrostatic surface properties, we employed various tools and techniques. Firstly, APBS^53^ was utilized to compute Poisson-Boltzmann electrostatics for each protein, using the corresponding PQR file. Nanoshaper^42^ was employed to generate a triangulated mesh representing the molecular surface. The charge at each vertex of the meshed surface was assigned using Multivalue, a tool within the APBS suite. Next, an in-house Python script was developed to map the electrostatic potential from the vertices to corresponding atoms. To obtain per residue values, we calculated the sum of all electrostatic potentials, as well as all the sum of all the positive or negative values. This process yielded three values: APBS_neg, APBS_pos, and APBS_sum for each residue.

We calculated electrostatic descriptors at the Fab, Fv, and CDR levels by integrating the residue-level electrostatic potentials (APBS_neg, APBS_pos, and APBS_sum) across the respective regions. For numbering antibody amino-acid variable domain sequences and the CDR regions we employed ANARCI^88^ using the Kabat scheme. This integration allowed us to derive the following electrostatic descriptors: Fab_APBS_pos, Fab_ APBS_neg, Fab_APBS_sum, Fv_APBS_pos, Fv_APBS_neg, Fv_APBS_sum, CDR_APBS_pos, CDR_APBS_neg, and CDR_APBS_sum.

### HPATCH: Hydrophobic Patch Calculations

To calculate the surface area of the given antibody structure, we developed a tool called HPATCH. HPATCH utilizes the PDB structure of an antibody to generate a triangulated solvent-excluded molecular surface (SES) using the Nanoshaper molecular surface.^42^ The SES construction a probe size of 1.5 Å and a density of 5 points per Å^²^. Each vertex on the surface is assigned a hydrophobicity scalar value based on four different scales: Kyte-Doolittle (KD), Wimley-White (WW), Eisenberg (EI), and Black & Mould (BM). These hydrophobicity values are scaled to a range between -5 (hydrophilic) and +5 (most hydrophobic). To account for clustering effects, the hydrophobicity value for each vertex is determined by calculating the average of all neighboring vertices within a Euclidean cutoff distance of 10 Å. For computational efficiency, we assume two surface vertices are neighbors if their associated atoms are within the same cutoff distance, thus only requiring a search for distances between associated atoms to identify neighbors. To further enhance computational speed, the code partitions the space into cubic boxes of size (2 x cutoff) and stores the atoms within each box. The algorithm then iterates over each atom and searches within the neighboring cubic boxes. Finally, the hydrophobicity values of the vertices are summed and assigned to their corresponding atoms. These values are further summed for all the atoms within each residue to obtain a residue-level hydrophobicity value. The hydrophobicity of each domain (CDR, Fv, Fab) is calculated by summing up the hydrophobicity values of all the corresponding residues that have a positive (hydrophobic) value.

### TAP descriptors

Therapeutic Antibody Profiler (TAP)^1^ descriptors for clinical antibodies were collected from Ref.,^65^ which were used in the analysis of PSR, Heparin, PK clearance and HIC. For the other sets (SEC set from Genetech and HMW data from Highland Games), we calculated the descriptors using the standard parameters (Fv built using AB2 and run at pH 7.4). We utilized a containerized version of the software licensed from OxPig.

### MOE protein properties

The MOE descriptors for clinical antibodies were collected from Ref.,^65^ which were used in the analysis of PSR, Heparin, PK clearance and HIC. For the remaining sets (SEC and HMW data), we used the following protocol to generate the descriptors. The software Molecular Operating Environment (MOE)^31^ from Chemical Computing Group was used to build the homology models of fragment antigen-binding (Fab) domains. The force field was set to the default Amber10 EHT using an internal and external dielectric values of 4 and 80, the non-bonded cutoff distances of 10 and 12 Å, and the Born solvation model. The prepared structures were energy minimized to root mean square gradient (RMSG) below 0.00001 kcal/mol/Å2. Protein properties for each Fab region were calculated using the Ensemble pH sampling module available in MOE, spanning two pH ranges, 4.5 to 6.5 6.4 to 8.4 (targeting pH=7.4). The number of conformations per unit of pH range was set to 25. The protein properties were calculated as Boltzmann weighted averages of the individual values from an ensemble of 50 total conformations.

### Additional *in silico* Descriptors

SCM and PfAbNet viscosity predictions were pulled from predictions provided by Apgar et al.^56^

### Accuracy Metrics

ROC-AUC is calculated using scikit-learn’s roc_auc_score method.^89^ As our predictions are not categorical, we treat them as probability estimates; the scikit-learn method then applies a moving threshold to achieve the ROC curve. Pearson *ρ*, which measures the linear correlation between two datasets, and Spearman *rho*, which measures the monotonic relationship between two datasets, are calculated using SciPy.^90^ Pearson *R*^2^ is the square of Pearson *rho* and is used in situations where magnitude of correlation is important, rather than direction. The coefficient of determination, *R*^2^, is calculated using scikit-learn’s r2_score method.^89^ As Pearson *R*^2^ and the coefficient of determination, *R*^2^, are not the exact same method, we stylize the former always as “Pearson *R*^2^.”

### *in vitro* and *in vivo* Assays

Aggregation state of the concentrated antibody samples were determined by Size Exclusion Chromatography using a protocol described previously.^91^ All other empirical data are from published datasets. The method details can be found in the corresponding publications. Viscosity data (PDGF38) was taken from Apgar et al. 2020.^62^ PK clearance data was taken from Grinshpun et al. 2021.^63^ PSR data was taken from Jain et al. 2017.^73^ Heparin data was taken from Kraft et al. 2020.^64^ Aggregation (%HWM/day) data was taken from Highland Games 2020.^34^ HIC data was taken from Jain et al. 2017.^40^

### Sequence Data

A set of 674 Clinical Stage Therapeutic (CST) sequences were downloaded from from Thera-SAbDab^77^ in August 2020. Due to structure prediction or simulation errors, we studied a subset of 557 sequences in the sections on sensitivity to structure model and the impact of conformational sampling.

## Acknowledgement

The authors gratefully thank Bob Kelley, Jessie Zhao, Jonathan Zarzar, Trevor Swartz, Nandhini Rajagopal, Shrenik Mehta and Jasper Lin for their valuable comments and suggestions. The authors also thank Thomas Hoeffel, Joseph Lipscomb, Nevin Cheung, and George Markov for their assistance in running our simulations smoothly on our high performance computing cluster.

